# Dual-loop DogTag engineering enables modular antigen display on human vault nanoparticles

**DOI:** 10.64898/2026.11.27.769278

**Authors:** Ismail Cem Yilmaz, Muzaffer Yildirim, Elif Sena Ozkoc, Gokcenaz Tuluoglu, Nilay Ayyildiz, Harun Onler, Emre Mert Ipekoglu, Bilgi Gungor, Ihsan Gursel, Mayda Gursel

## Abstract

Self-assembling protein nanoparticles can enhance immune responses by displaying multiple copies of an antigen on their surface. Most previous studies using vault nanoparticles have focused on loading proteins inside the particle or modifying its ends. Here, we introduced two DogTag peptides into separate surface-exposed loops of human major vault protein (MVP), creating up to 156 potential attachment sites on each 78-subunit vault. The engineered MVP formed vault-sized particles in suspension HEK cells, to which separately produced, stabilized influenza H5N1 Clade 2.3.4.4b hemagglutinin (HA) fused to DogCatcher was covalently attached. By varying the amount of HA added, we obtained preparations with estimated loadings of approximately 32, 64, and 124 HA trimers per vault. Structural models showed that increasing HA loading reduced the median distance between neighboring HA trimers from 13.3 to 8.8 nm, while electron microscopy showed that highly loaded particles retained their characteristic vault morphology. Following subcutaneous priming and intranasal boosting in mice, highly loaded HA-decorated vaults induced significantly higher HA-specific serum IgG, IgG1, IgG2a, and IgA titers than soluble HA. Covalent attachment of HA to vaults also resulted in significantly higher IgG2a titers than the non-conjugated mixture of vaults and HA, reaching levels similar to those induced by CpG ODN adjuvanted HA. Despite the strong HA-specific antibody responses, anti-vault IgG titers remained low. Our approach adds high-density, adjustable antigen display to the established cargo-loading and targeting capabilities of human vaults, offering the possibility of combining all three functions in a single vaccine particle.

## Introduction

Vaults are among the largest ribonucleoprotein particles found in eukaryotic cells. The human vault shell is composed of 78 copies of the ∼100-kDa major vault protein (MVP), arranged as two antiparallel half-shells of 39 subunits each (Kedersha and Rome, 1986; Tanaka et al., 2009; Lapenta et al., 2026). Native mammalian vaults also contain the associated proteins TEP1 and PARP4 together with small vault RNAs, whereas MVP alone forms the structural shell and is sufficient for recombinant particle assembly in the absence of other native components. Assembled vaults adopt a characteristic barrel-like geometry measuring approximately 40 × 40 × 66 nm, providing a large exterior surface and a spacious internal cavity (Tanaka et al., 2009; Lapenta et al., 2026). Their nanoscale dimensions, repetitive subunit architecture, and accessible exterior and interior compartments make recombinant vaults well suited for multivalent antigen display and cargo encapsulation.

Compared with the 78-subunit vault shell, most established protein nanoparticle vaccines rely on smaller self-assembling scaffolds, such as the 24-subunit ferritin cage (∼12 nm) and the 60-subunit lumazine synthase particle (Jardine et al., 2013; Mastrangelo and Di Pisa, 2026). Genetic fusion of viral glycoproteins, including influenza hemagglutinin (HA), to ferritin generates nanoparticles displaying eight HA trimers per particle and elicits strong antibody responses across multiple vaccine studies (Kanekiyo et al., 2013; Yassine et al., 2015; Houser et al., 2022). More recently, SpyTag/SpyCatcher chemistry has enabled separately produced antigens to be covalently attached to pre-assembled ferritin and lumazine synthase scaffolds, avoiding the need to express the antigen and scaffold as a single fusion protein (Zhang et al., 2020).

In contrast to these scaffold-based antigen display platforms, recombinant vault engineering has focused primarily on encapsulating cargo inside the particle lumen or modifying the terminal ends of MVP. Fusing proteins to the MVP-interaction domain (INT) from PARP4/VPARP directs them into the vault interior, an approach used to package enzymes and cytokines (Kickhoefer et al., 2005; Lou et al., 2026) as well as Chlamydia antigens for intranasal vaccination in mice (Champion et al., 2009; Jiang et al., 2017). Previous examples of exterior targeting include C-terminal fusions of targeting peptides or protein ligands to MVP at the particle caps (Kickhoefer et al., 2009).

Controlling surface antigen density and spacing is important for vaccine design because multivalent geometry influences B-cell receptor clustering, germinal-center recruitment, and antibody maturation, though optimal spacing varies across antigens and B-cell repertoires (Bachmann et al., 1993; Kato et al., 2020; Veneziano et al., 2020; Wamhoff et al., 2024). Because the elongated vault shell provides a large exterior surface, it offers substantial space for arranging bulky antigens across a wide density range. Expressing the antigen separately and conjugating it after particle assembly also avoids the need to produce large viral glycoproteins as direct fusions to nanoparticle subunits, which can impose constraints on protein folding and particle assembly (Nguyen and Tolia, 2021; Zhang et al., 2020). The DogTag/DogCatcher system is well suited to this modular strategy because DogTag was developed specifically for insertion into internal protein loops and forms a spontaneous covalent isopeptide bond with DogCatcher (Keeble et al., 2022). Introducing DogTag into surface-exposed loops of MVP therefore enables post-assembly covalent attachment across the vault exterior with tunable antigen loading.

Here, we engineered human MVP to carry two DogTag peptides on separate surface-exposed loops, generating up to 156 potential attachment sites across the 78-subunit vault shell. We produced and purified these dual-DogTag vaults, covalently conjugated separately expressed H5N1 Clade 2.3.4.4b HA–DogCatcher across a broad titration range, and quantified antigen loading biochemically. We then used structural modelling and negative-stain scanning transmission electron microscopy (STEM) to assess antigen organization and particle morphology. Finally, we evaluated the in vivo immunogenicity of HA-decorated vaults in mice following a subcutaneous prime and intranasal boost, assessing both HA-specific antibody responses across immunoglobulin classes and reactivity against the vault scaffold. Together, these findings establish dual-loop MVP engineering as a modular platform for tunable multivalent antigen display on human vault nanoparticles.

## Results

### Dual-loop DogTag insertion into MVP creates 156 potential attachment sites arranged in four tiers around the vault body

To place DogTag sites on the outside of the vault, we inserted DogTag into two surface-exposed loops of human major vault protein (MVP) (NP_005106.2). Using the human vault structure (PDB 9QW9) as a guide, we selected the D131–K136 loop in repeat domain R3 and the D345–K348 loop in R7 (Fig. 1A, B). Each loop is present once in every MVP subunit and, in the assembled 78-subunit vault, the two sites form four circumferential tiers around the vault body.

**Figure 1.**
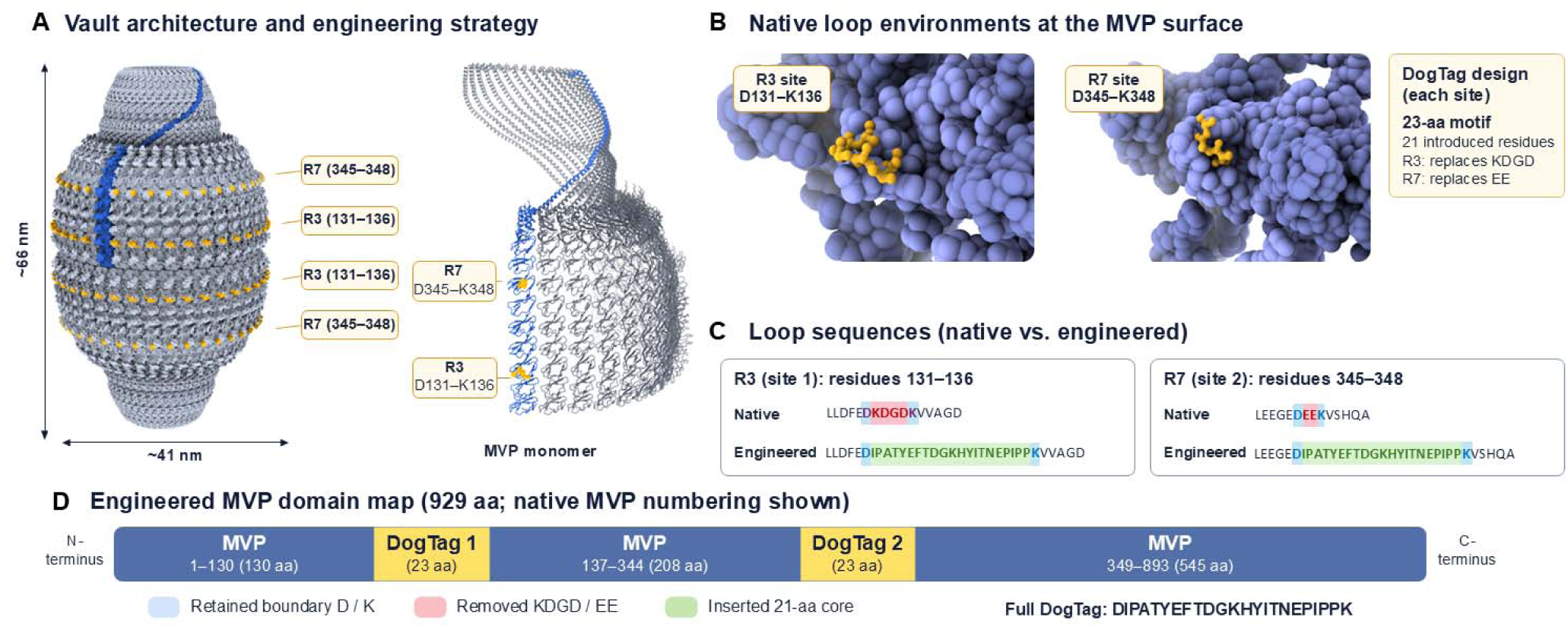
Dual-loop DogTag engineering of human major vault protein. **(A)** Architecture of the human vault (PDB 9QW9) showing a representative MVP protomer in blue and the two target loop positions (R3 and R7) in gold. Each of the 78 MVP subunits contributes one R3 and one R7 site, providing 156 potential attachment sites arranged in four circumferential tiers. The positions of the R3 (D131–K136) and R7 (D345–K348) loops are also indicated on the MVP protomer. **(B)** Structural environments of the two native target loops, with the corresponding residues shown in gold against the surrounding MVP surface. **(C)** Native and engineered sequences at the two DogTag insertion sites. At R3, residues 132–135 (KDGD) were replaced, whereas at R7, residues 346–347 (EE) were replaced by the 21 internal residues of DogTag. The native flanking Asp and Lys residues were retained at both sites to form the complete 23-residue DogTag motif (DIPATYEFTDGKHYITNEPIPPK). **(D)** Linear map of the engineered MVP showing the positions of the two DogTag motifs. The modifications resulted in net additions of 17 and 19 residues at R3 and R7, respectively, increasing the length of MVP from 893 to 929 amino acids. Residue numbering refers to the native human MVP sequence (NP_005106.2).

The insertions were designed to retain the native Asp and Lys residues at the ends of each loop, which also form the terminal residues of the 23-residue DogTag sequence (Fig. 1C). At the R3 site, residues 132–135 (KDGD) were replaced while D131 and K136 were retained. At the R7 site, residues 346–347 (EE) were replaced while D345 and K348 were retained. This introduced 21 new residues at each site, giving a net increase of 17 residues at R3 and 19 residues at R7. The engineered MVP therefore contains 929 amino acids compared with 893 amino acids in native MVP (Fig. 1D). Since every MVP subunit carries both DogTag insertions, a vault assembled from 78 engineered MVP subunits contains up to 156 potential DogTag attachment sites.

### Expression and purification of dual-DogTag vault nanoparticles in suspension HEK cells

To determine whether the dual-loop modifications were compatible with production of vault particles, the engineered MVP was expressed in suspension HEK cells and purified by sequential Capto Core 700 and qEV1 size-exclusion chromatography (Fig. 2A). Stain-free SDS–PAGE showed a prominent band at the expected migration position of engineered MVP (∼120 kDa) in the transfected preparation, which was retained through Capto Core purification and was evident in qEV fractions 6 and 7 (Fig. 2B). In contrast, matched mock qEV fractions lacked a comparable band at this position.

**Figure 2.**
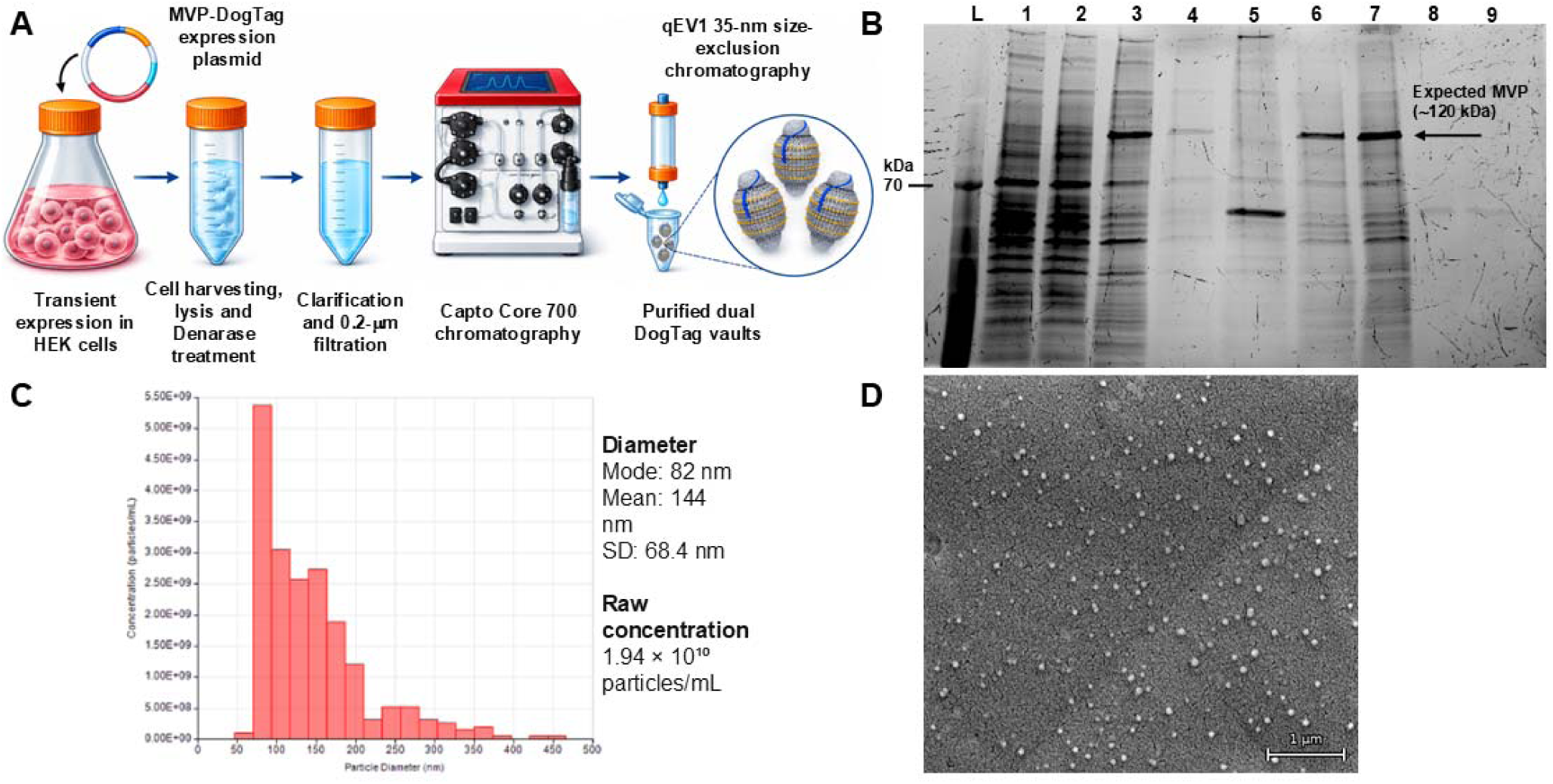
Production and recovery of engineered dual-DogTag vault particles. **(A)** Schematic of the production and purification workflow. Dual-DogTag human MVP was expressed in suspension HEK cells for four days following plasmid transfection. Cell lysates were treated with Denarase, clarified by 0.2-µm filtration and sequentially purified by Capto Core 700 chromatography and qEV1 35-nm size-exclusion chromatography. **(B)** Stain-free SDS–PAGE of samples collected during purification. L, molecular-weight ladder; lane 1, mock lysate; lane 2, transfected lysate; lanes 3–4, transfected Capto Core 700 peaks 1–2; lane 5, mock Capto Core 700 peak 1; lanes 6–7, transfected qEV fractions 6–7; lanes 8–9, matched mock qEV fractions 6–7. The 70-kDa ladder position is indicated. The arrow marks the expected migration position of engineered MVP (∼120 kDa). **(C)** Particle size distribution and concentration of the post-qEV preparation determined by tunable resistive pulse sensing (TRPS). **(D)** Representative scanning electron micrograph of the post-qEV preparation. Scale bar, 1 µm.

To confirm that the purified MVP assembled into intact nanoparticles rather than remaining as free protein subunits, we measured particle size and concentration by tunable resistive pulse sensing (TRPS) and examined the samples by scanning electron microscopy (SEM). TRPS analysis of the post-qEV preparation detected a particle population with a modal diameter of 82 nm, a mean diameter of 144 nm (S.D, 68.4 nm), and a concentration of 1.94x10^10^ particles/mL (Fig. 2C). SEM imaging of the same preparation confirmed the presence of multiple particles with sizes and morphology consistent with vault particles (Fig. 2D).

Together, these results show that inserting two DogTag loops into human MVP does not prevent particle assembly, allowing intact vault-sized particles to be produced and purified from suspension HEK cells.

### HA–DogCatcher can be covalently attached to dual-DogTag vaults over a broad loading range

To test whether the engineered vaults can bind external antigens, we produced a stabilized influenza HA–DogCatcher fusion protein containing a GSAS replacement at the furin-cleavage site, three stabilizing substitutions (H371W, K396I, and E448I) (Milder et al., 2022), a C-terminal T4 foldon trimerization domain, and DogCatcher (Fig. 3A). HA was chosen as a large trimeric viral glycoprotein to test whether the engineered vault could carry a bulky antigen. This antigen was expressed and purified from suspension HEK cells separately from the vault scaffold and later combined with engineered vaults for covalent conjugation to the surface.

**Figure 3.**
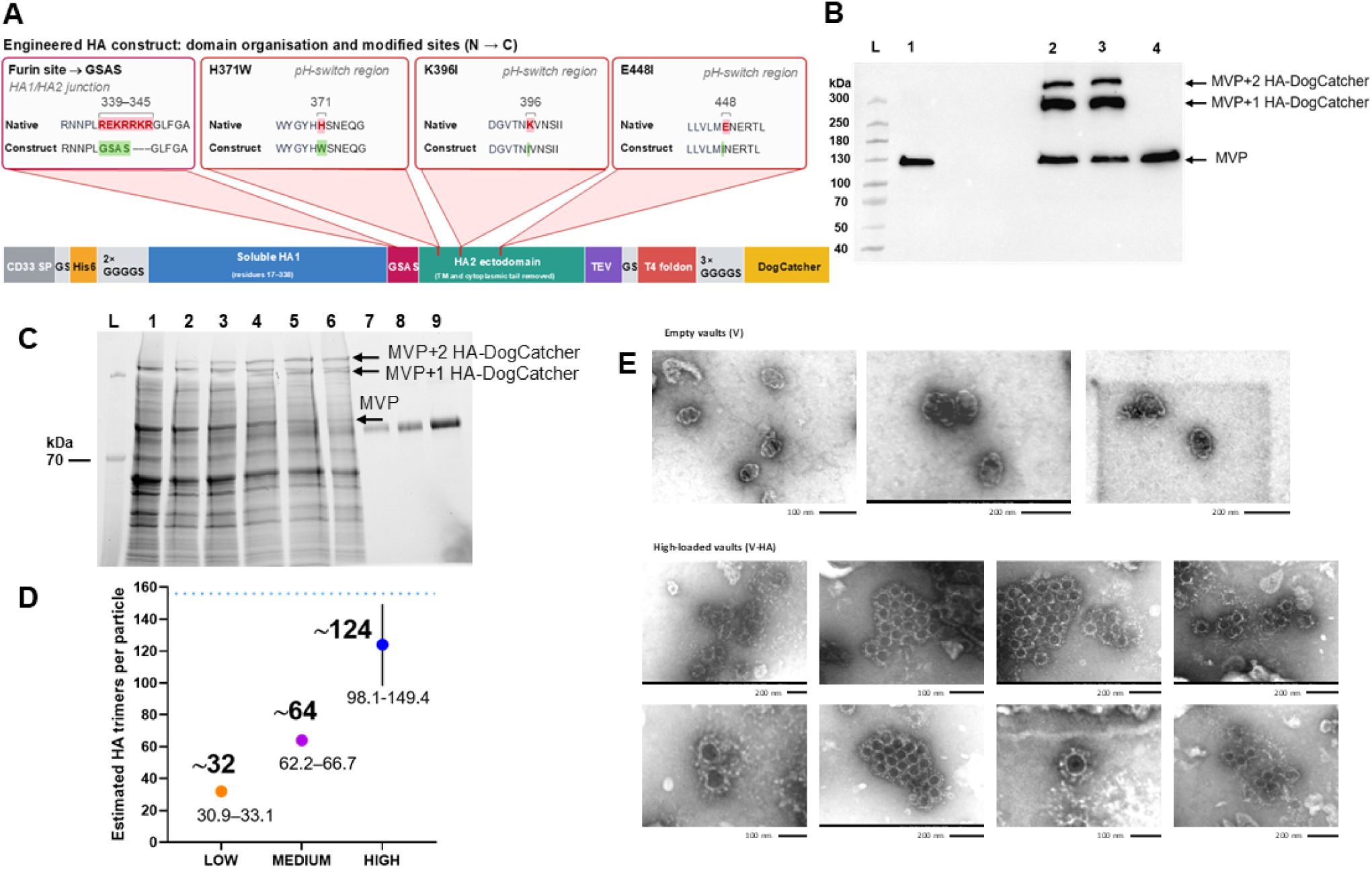
Covalent loading of stabilized HA–DogCatcher onto dual-DogTag vault particles. **(A)** Schematic of the HA–DogCatcher construct used for vault conjugation. The soluble HA ectodomain contains a GSAS substitution at the native furin-cleavage site and three stabilizing substitutions (H371W, K396I and E448I). The construct includes an N-terminal CD33 signal peptide, His₆ tag, (GGGGS)₂ linker, HA1 and HA2 ectodomain regions, TEV protease cleavage site, T4 fibritin foldon trimerization domain, (GGGGS)₃ linker and C-terminal DogCatcher. **(B)** Anti-MVP immunoblot of vault–HA conjugation. L, molecular-weight ladder; lane 1, dual-DogTag vaults alone; lanes 2–3, vaults incubated with HA–DogCatcher; lane 4, vaults incubated with the non-cognate HA–SpyTag control. The positions of unmodified MVP and MVP carrying one or two HA–DogCatchers polypeptides are indicated. **(C)** Stain-free SDS–PAGE of vault preparations incubated with increasing amounts of HA–DogCatcher. L, molecular-weight ladder; lane 1, vault-only preparation; lanes 2–5, vault preparations incubated with 25, 50, 100 and 200 µg HA–DogCatcher, respectively; lane 6, separate high-loading preparation generated with 800 µg HA–DogCatcher; lanes 7–9, purified HA–DogCatcher standards containing 0.5, 1.0 and 2.0 µg protein per lane, respectively. **(D)** Estimated HA loading in qEV-purified vault preparations. Particle-associated HA was quantified by anti-His-tag ELISA and normalized to vault-sized particle counts determined by TRPS after excluding larger-particle populations. The preparations generated with 50, 100 and 800 µg HA–DogCatcher define the LOW, MEDIUM and HIGH loading states, respectively. Values represent estimated HA trimer equivalents per particle, with the corresponding estimation ranges indicated. The dotted line indicates the 156 theoretical DogTag attachment sites per vault. **(E)** Representative negative-stain scanning transmission electron microscopy (STEM) images of empty (V) and highly loaded HA-decorated vaults (V-HA). Scale bars are as indicated.

To verify that attachment depended on specific DogTag–DogCatcher covalent coupling, we incubated dual-DogTag vaults with HA–DogCatcher or with a non-cognate HA–SpyTag control and analyzed the reactions by anti-MVP immunoblotting. Vaults incubated with HA–DogCatcher showed two higher-molecular-weight bands above the unmodified ∼120 kDa MVP band, corresponding to MVP chains covalently linked to one or two HA–DogCatcher polypeptides (Fig. 3B). By contrast, no shifted bands appeared with the non-cognate HA–SpyTag control, confirming specific covalent conjugation through the engineered DogTag sites.

To determine whether antigen loading could be controlled on the same scaffold, we incubated a fixed amount of vaults with increasing amounts of HA–DogCatcher (25, 50, 100, and 200 µg), purified them via qEV size-exclusion chromatography to remove unconjugated HA trimers and analyzed reactions by Stain-Free SDS–PAGE. Adding more HA–DogCatcher progressively reduced the unmodified MVP band while increasing the higher-molecular-weight conjugated bands (Fig. 3C). Across this controlled loading series, the apparent loss of unmodified MVP rose from 3.3% at 25 µg HA to 27.2%, 44.5%, and 67.2% at 50, 100, and 200 µg, respectively (Supplementary Table 1A1). A separate maximal-loading preparation made with 800 µg HA under different reaction conditions, in which HA was added to lyophilized vaults to compensate for the increase in volume upon mixing, showed an 82.9% reduction in unmodified MVP and was used as a separate high-loading reference. Purified HA–DogCatcher standards run on the same gel showed a linear Stain-Free response over the 0.5–2.0 µg range (R² = 0.997), indicating that the HA bands were not saturated under these conditions (Supplementary Table 1A3).

To estimate how many DogTag sites reacted per vault, we quantified the unmodified MVP, MVP+1 HA–DogCatcher and MVP+2 HA–DogCatcher bands and corrected the shifted-band signals for their different tryptophan contents. Engineered MVP contains 7 tryptophan residues and HA–DogCatcher contains 11, giving 18 and 29 tryptophan residues for the singly and doubly conjugated MVP species, respectively. Combining the loss of unmodified MVP with the tryptophan-corrected distribution of the two shifted species gave approximately 32, 54, and 81 reacted DogTag sites per vault for the 50, 100, and 200 µg HA inputs, respectively (Supplementary Table 1A2). The 25 µg lane was too weak for reliable analysis of the shifted species, and the 800 µg sample was treated only semiquantitatively because it was prepared under different reaction conditions.

Next, to estimate the average number of HA trimers per vault after removing unbound antigen, we measured the HA concentration in the qEV-purified particle fractions by anti-His-tag ELISA and normalized the resulting HA trimer count to the number of vault-sized particles measured by TRPS, excluding larger-particle populations (Fig. 3D). The 25–200 µg HA series gave approximately 3.5 × 10¹¹ vault-sized particles per pooled preparation. Using this denominator, the +50 µg preparation yielded 30.9–33.1 particle-associated HA trimers per vault and defined the LOW state (∼32), while the +100 µg preparation yielded 62.2–66.7 and defined the MEDIUM state (∼64). The +200 µg preparation yielded 73.1–78.3 HA trimers per particle. The separate maximal-loading preparation yielded 98.1–149.4 HA trimers per vault (midpoint, 123.8) and defined the HIGH state (∼124). The ELISA and TRPS values used for these particle-normalized loading estimates are provided in Supplementary Table 1B.

Although the ELISA/TRPS method estimated the average number of HA trimers per vault whereas gel densitometry estimated reacted DogTag sites, the two approaches gave values on a similar scale (∼32, ∼64 and ∼76 HA trimers versus ∼32, ∼54 and ∼81 reacted DogTag sites across the 50, 100 and 200 µg inputs, respectively). The two assays measure different quantities and are consistent with an overall increase in loading across the controlled series.

Then, to see how HA attachment affected the physical appearance of the particles, we imaged empty and highly loaded vaults by negative-stain scanning transmission electron microscopy (STEM; Fig. 3E). Both preparations contained vault-sized particles, supporting retention of vault-like morphology after conjugation. In the highly loaded preparation, near-end-on views frequently showed a dark, stain-filled central opening whose appearance and approximate dimensions were consistent with the hollow central lumen reported for native vaults (Tanaka et al., 2009; Lapenta et al., 2026), further supporting preservation of the characteristic barrel architecture after HA conjugation. Whereas empty vaults showed smooth, undecorated edges, highly loaded vaults were frequently surrounded by irregular peripheral density consistent with surface-bound HA.

In summary, these results show that dual-DogTag vaults can be covalently coated with separately produced HA–DogCatcher across a wide, controllable range of loading levels without redesigning the particle. Gel densitometry and ELISA/TRPS provided independent estimates of increasing HA loading, while STEM imaging supported retention of vault-like morphology at high loading. These measurements defined the experimental LOW (∼32), MEDIUM (∼64), and HIGH (∼124) loading states used in the structural modelling below.

### Structural modelling of HA loading on vaults

In order to determine whether the experimentally estimated HA copy numbers can be accommodated on the vault surface without steric clashes and to assess the effect of loading on antigen spacing, we generated structural models of the three representative states: LOW (32 HA trimers per vault), MEDIUM (64 trimers), and HIGH (124 trimers) (Fig. 4A). For the models, the HA–foldon core was treated as a rigid steric volume that could not overlap with the vault shell or neighboring HA trimers, while the DogTag–DogCatcher–linker segment was free to assume flexible conformations. Representative configurations for all three loading states satisfied the applied steric criteria for the modeled rigid antigen cores. Higher HA occupancy reduced the three-dimensional nearest-neighbour distance between trimers, with median spacing falling from 13.3 nm at LOW loading (range, 8.7–23.5 nm) to 10.2 nm at MEDIUM loading (8.4–16.0 nm) and 8.8 nm at HIGH loading (7.9–10.7 nm) (Fig. 4A). Thus, the models predict loading-dependent changes in local antigen spacing on the same vault scaffold.

**Figure 4.**
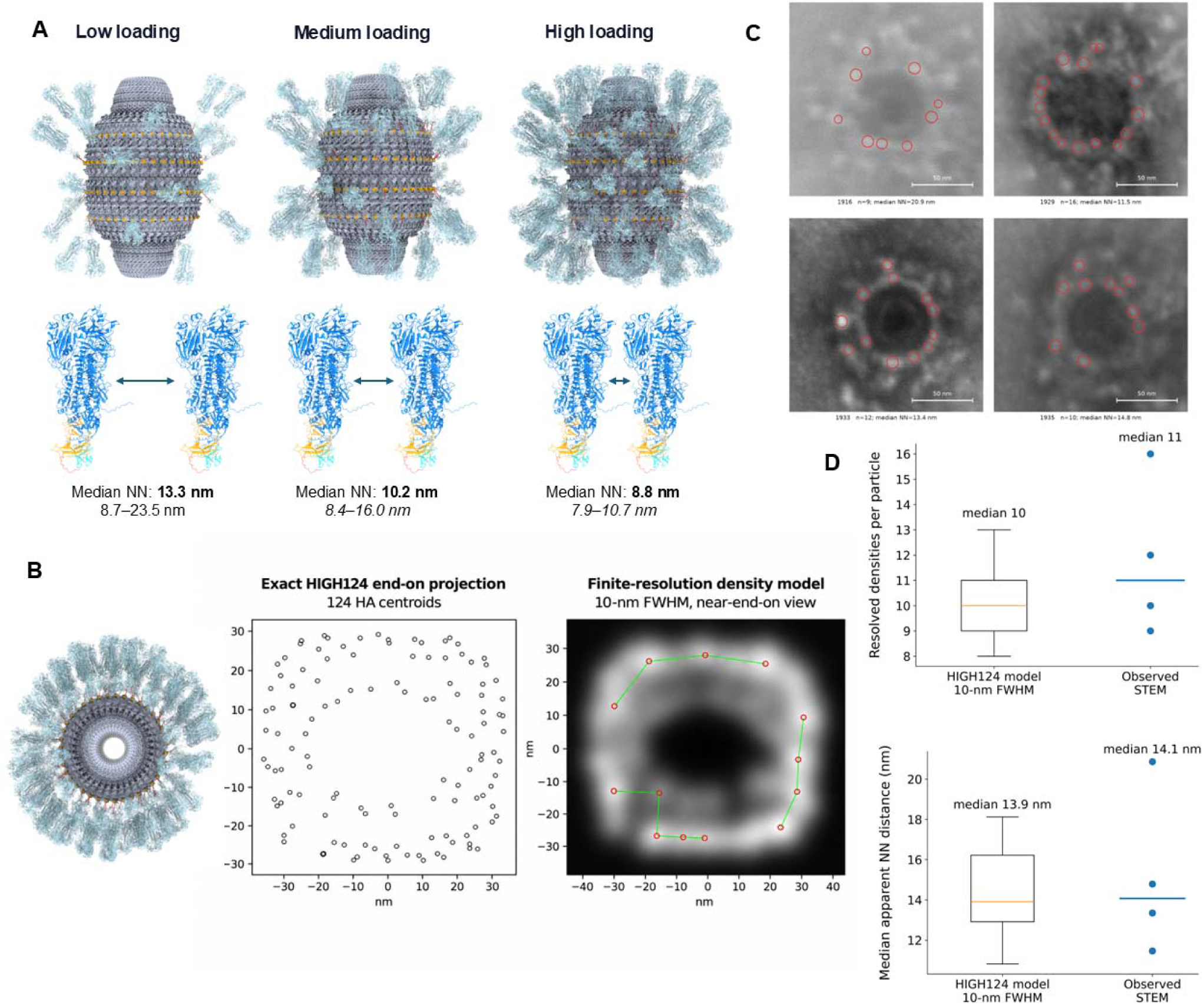
Structural modelling of HA organization on vault particles and comparison with STEM images. **(A)** Structural models of vaults carrying 32 (LOW), 64 (MEDIUM) and 124 (HIGH) HA trimers, based on the experimentally estimated loading values. The median three-dimensional nearest-neighbour (NN) distances between HA trimers are shown below each model. **(B)** End-on view of the HIGH124 model (left), showing the projected positions of the 124 HA trimer centres (middle) and their simulated appearance at an assumed spatial resolution of 10 nm (right). **(C)** Negative-stain STEM images of four highly loaded vault particles viewed end-on. Red circles mark the individual spots visible around the particle surface, which were used for counting and distance measurements. Scale bars, 50 nm. **(D)** Comparison of the number of visible spots per particle (upper panel) and their median NN distances (lower panel) in 37 simulated views and four STEM images.

We next compared the HIGH (124-trimer) model with negative-stain STEM images of the maximally loaded preparation. Because the highly loaded vaults adopted predominantly near-end-on orientations on the grid and negative-stain STEM yielded 2D projections with limited spatial resolution across multiple attachment sites (156 nominal DogTag positions), direct 3D reconstruction from the images was not feasible. Therefore, in order to see what a vault carrying 124 HA trimers should look like from an end-on angle, we projected our 3D model onto a flat 2D plane and applied an assumed 10-nm effective image-formation parameter (Fig. 4B). In this end-on projection, HA trimers sitting in four rings stacked along the height of the vault lined up behind one another and merged into single spots, producing far fewer visible spots than the 124 modeled trimers. Next, to check whether this visual overlap explains our microscopy results, we counted the distinct spots around the outer edge and measured their spacing in four isolated, end-on STEM particles (Fig. 4C). These particles showed 9, 16, 12, and 10 spots around the perimeter, giving a median of 11 spots per particle and a median spacing of 14.1 nm between spots (range, 11.5–20.9 nm). To compare these experimental numbers directly against our model, we performed the same measurement across 37 simulated end-on views of the 124-HA vault, which yielded a median of 10 spots spaced 13.9 nm apart (Fig. 4D). The descriptive agreement between simulated views (10 spots, 13.9 nm) and the four STEM particles (11 spots, 14.1 nm) is compatible with the HIGH model.

Overall, the models support the geometric feasibility of the representative high-loading state under the stated assumptions and predict that changing average loading alters antigen spacing.

### HA-decorated vaults enhance antigen-specific antibody responses in mice

To evaluate the immunogenicity of HA-decorated vaults *in vivo*, 6–8-week-old female BALB/c mice (n = 8 per group) were immunized with a subcutaneous (SC) prime followed by an intranasal (IN) boost. The five experimental groups received empty vaults (Vault), soluble HA (HA), a mixture of empty vaults and soluble HA (Vault+HA), HA-decorated vaults (Vault-HA), or HA formulated with CpG ODN (HA+CpG ODN). All HA-containing groups received 2 µg HA at priming and 1 µg HA at boosting. For the Vault-HA group, we used the highly loaded preparation generated with 800 µg HA–DogCatcher (Fig. 3C, lane 6), from which unconjugated HA was removed by qEV size-exclusion chromatography. Vault-containing groups received approximately 3.3 × 10^10^ particles at priming and 1.65 × 10^10^ particles at boosting. HA-specific serum IgG, IgG1, IgG2a and IgA endpoint titers were determined by ELISA 10 days after the booster injection (Fig. 5A).

**Figure 5.**
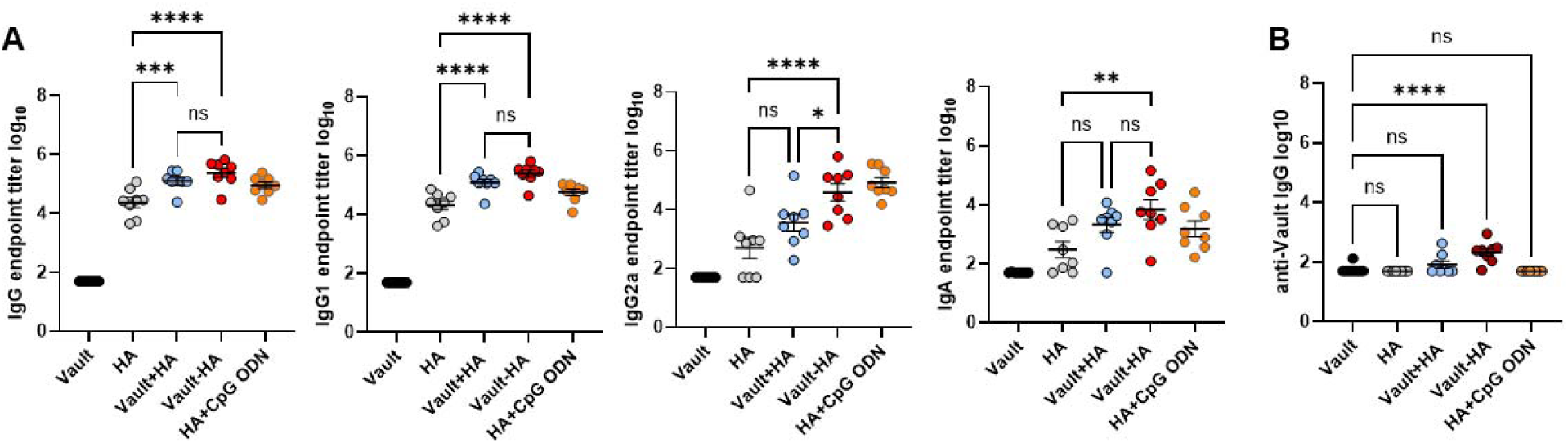
Humoral immune responses induced by HA-decorated vaults in mice. Mice (n= 8 per group) were primed subcutaneously (2 µg HA) and boosted intranasally (1 µg HA) with empty vaults (Vault), soluble HA (HA), a physical mixture of empty vaults and HA (Vault+HA), qEV-purified HA-decorated vaults (Vault-HA), or HA adjuvanted with CpG ODN (HA+CpG ODN). Serum samples collected 10 days after boosting were analyzed by ELISA. (A) HA-specific serum total IgG, IgG1, IgG2a, and IgA log_10_ endpoint titers are shown. (B) Anti-vault serum IgG log_10_ endpoint titers measured on vault-coated plates. Statistical significance was determined by one-way ANOVA followed by Šídák’s multiple-comparisons test (*p < 0.05,**p < 0.01, ***p < 0.001, ****p < 0.0001; ns, not significant).

As expected, mice that received empty vaults showed no detectable HA-specific antibody responses. In contrast, Vault-HA significantly increased total IgG (p < 0.0001), IgG1 (p < 0.0001), IgG2a (p < 0.0001) and IgA (p < 0.01) titers compared with soluble HA. Mixing empty vaults with soluble HA also significantly increased total IgG (p < 0.001) and IgG1 (p < 0.0001) titers, indicating an adjuvant-like effect of the mixed formulation under these conditions. However, Vault-HA elicited significantly higher IgG2a titers than Vault+HA (p < 0.05), with titers in the range observed for HA adjuvanted with CpG ODN. Furthermore, Vault-HA was the only vault-containing formulation that significantly increased serum IgA titers relative to soluble HA (p<0.01).

In an independent preliminary experiment (n = 5 mice per group), mice received the same HA doses and SC–IN immunization regimen, but with a Vault-HA preparation from which unconjugated HA had not been removed and at lower particle doses (1–2 × 10⁸ vaults at priming and 7 × 10⁸ at boosting). Despite these differences, Vault-HA elicited significantly higher total IgG (p < 0.05), IgG1 (p < 0.05) and IgA (p < 0.0001) titers than Vault+HA as well as significantly higher IgG1 (p<0.05) titers than soluble HA, whereas IgG2a titers did not differ significantly between these three groups (Fig. S1A). Collectively, these findings show that HA-decorated vaults enhance HA-specific serum IgG and IgA responses without an added adjuvant, with broader enhancement across antibody classes in the main experiment.

We also examined whether immunization induced antibodies against the vault scaffold by measuring serum IgG reactivity against vault-coated ELISA plates in both experiments (Figure 5B and Fig. S1B). Despite the substantially higher number of vault particles administered in the main experiment (165–330-fold at priming and 24-fold at boosting), vault-reactive IgG titers remained low across all groups (Figure 5B and Fig. S1B). Mice that received soluble HA, Vault+HA or HA+CpG ODN showed little or no reactivity against vault-coated plates and did not differ significantly from mice immunized with empty vaults alone. Only mice immunized with Vault-HA showed a significant increase in vault-reactive IgG titers compared with mice vaccinated with empty vaults (p < 0.0001), although these titers were approximately three orders of magnitude lower than the corresponding HA-specific IgG titers (Fig. 5A, B). Similarly, in the preliminary cohort immunized with lower vault particle numbers, anti-vault IgG titers remained low across all groups, with only minor reactivity detected in mice that received Vault-HA (p<0.05) or HA+CpG ODN (p<0.05) with respect to empty vaults (Fig.S1B). Taken together, the results of both immunization experiments are consistent with low vault-reactive serum IgG relative to HA-specific IgG under the conditions tested.

## Discussion

Recent high-resolution structures of human vaults show that 78 copies of the major vault protein (MVP) assemble into a large, highly ordered shell with an extensive external surface and a large internal cavity (Lapenta et al., 2026). To exploit this repetitive surface for multivalent antigen display, we engineered human MVP to carry two DogTag sites in separate surface-exposed loops, creating up to 156 potential attachment sites per particle. DogTag was particularly suitable for this approach because it was designed for insertion into protein loops and enables covalent attachment of DogCatcher-fused proteins (Keeble et al., 2022). Our results show that the vault architecture tolerates these modifications without preventing particle assembly and allows the attachment of a large trimeric viral antigen after assembly.

A key advantage of this design is that the antigen does not need to be genetically fused to MVP. Instead, vaults and antigens are produced separately and conjugated after assembly via DogTag–DogCatcher, allowing loading to be tuned simply by varying the input antigen ratio. In most self-assembling protein nanoparticles, antigen copy number and spacing are fixed by scaffold symmetry and/or genetic fusion design. Genetic fusion of influenza HA to ferritin displays eight HA trimers, while lumazine synthase and I53-50 similarly arrange antigens according to fixed particle geometries (Kanekiyo et al., 2013; Walls et al., 2020). Changing antigen density or spacing in those systems typically requires re-engineering the fusion, linker, or scaffold. Here, antigen density can be adjusted on the same pre-assembled vault: the LOW, MEDIUM, and HIGH preparations carried approximately 32, 64, and 124 HA trimers per particle, reducing the median nearest-neighbour distance from 13.3 nm to 10.2 nm and 8.8 nm in our structural models. This makes it straightforward to test different antigen densities when the optimal spacing is not known in advance. Varying loading also tests how much surface decoration the vault can accommodate before steric crowding limits conjugation or destabilizes the particle. In dual-functionalized ferritin nanocages, increasing the fraction of a bulky protein fusion reduced cage stability, requiring an intermediate ratio to preserve particle integrity (Shen et al., 2026). By contrast, our structural models accommodated the LOW, MEDIUM, and HIGH states without steric overlap, and highly loaded vaults maintained their characteristic morphology in STEM images.

The differences in antigen spacing produced by changing HA loading may also be biologically relevant. Antigen spacing can influence B-cell activation, but there does not appear to be a single spacing that is optimal for all antigens. Veneziano et al. reported stronger B-cell signalling when an HIV immunogen was displayed at wider spacing, reaching approximately 25–30 nm, whereas Ellis et al. found that decreasing the spacing of trimeric influenza HA-head antigens within the 10–12 nm range improved antibody potency and breadth (Veneziano et al., 2020; Ellis et al., 2023). These differences probably reflect the antigen, epitope accessibility, BCR affinity and the response being measured. A scaffold on which antigen spacing can be changed without rebuilding the particle may therefore be useful for testing different presentation densities.

Consistent with the advantages of dense trimer spacing reported for HA-conjugated nanoparticles (Ellis et al., 2023), in our study, displaying HA trimers at high density on the vault surface induced strong HA-specific IgG, IgG1, IgG2a and serum IgA responses without an added adjuvant. Although co-administration of empty vaults with soluble HA (Vault+HA) also enhanced IgG and IgG1 responses, covalent attachment of HA (Vault-HA) resulted in significantly higher IgG2a titers than the mixed formulation, reaching levels similar to those induced by HA adjuvanted with CpG ODN. Furthermore, Vault-HA was the only vault-containing formulation that significantly increased serum IgA titers compared with soluble HA. Whether antigen density and spacing on the vault surface influence these responses remains to be determined.

In addition to enhancing responses to the displayed antigen, an effective nanoparticle scaffold must also avoid provoking dominant anti-carrier immunity. Many protein based nanoparticle scaffolds use bacterial or computationally designed proteins that can trigger responses against the carrier rather than just the antigen. For example, in human clinical trials of the lumazine synthase-based eOD-GT8 60-mer vaccine, although HIV-specific B cells were induced, strong anti-carrier immunity also developed, including anti-lumazine synthase antibodies (Leggat et al., 2022) and lumazine-synthase-specific CD4 T cells in 93% of recipients after two doses (Cohen et al., 2023). Anti-scaffold antibodies have likewise been reported for ferritin, I53-50, and other protein nanoparticle platforms (Kanekiyo et al., 2013; Marcandalli et al., 2019; Kelly et al., 2020; Brouwer et al., 2021; Arunachalam et al., 2021; Tan et al., 2021). In contrast, the scaffold used in our study is derived from human MVP, a highly conserved mammalian protein. Repeated administration of recombinant vaults carrying Gaussia luciferase, mCherry, a chlamydial PmpG peptide or ovalbumin did not elicit detectable anti-MVP antibodies in mice, whereas vaults carrying the major outer membrane protein (MOMP) of Chlamydia did, suggesting that the nature of the associated antigen can influence scaffold immunogenicity (Champion et al., 2009; Jiang et al., 2017). Furthermore, no chronic inflammation or fibrosis was observed in major organs 60 days after immunization with empty or PmpG-containing vaults (Jiang et al., 2017). Consistent with these observations, mice immunized with empty vaults or Vault+HA in our study showed little or no vault-reactive IgG in either experiment, even when the particle dose was increased 165–330-fold at priming and 24-fold at boosting. Although HA-decorated vaults induced detectable vault-reactive IgG, these titers remained low compared with HA-specific IgG titers (Fig. 5B and Fig. S1B).

Beyond its relatively low intrinsic immunogenicity, the vault offers additional regions that can be engineered for different functions. The C-terminus of MVP, which is exposed at the two caps of the particle, has previously been modified with epidermal growth factor (EGF) or IgG-binding peptides to enable direct or antibody-mediated targeting of cell-surface receptors (Kickhoefer et al., 2009; Han et al., 2011; Tomaino et al., 2024). Thus, the cap region could hypothetically be used for cell-specific targeting while the DogTags on the side walls could serve as antigen attachment sites. Furthermore, the large internal cavity of the vault remains available for cargo loading. Fusion to the VPARP-derived INT domain has been used to package enzymes, fluorescent proteins, vaccine antigens and immunomodulatory proteins within recombinant vaults (Kickhoefer et al., 2005; Champion et al., 2009; Kar et al., 2011; Jiang et al., 2017). Recent structural work has also shown that recombinant human vaults can accommodate several hundred molecules of IL-2 per particle (Lou et al., 2026). Thus, a potential advantage of vaults over other platforms, such as ferritin, lumazine synthase and designed icosahedral cages, lies in the possibility of combining antigen display, cell-specific targeting and cargo delivery within a single particle. Our work complements the existing targeting and cargo-loading strategies by introducing a means of covalent antigen display on the outer surface of the vault.

The surface-display approach we employed may not be limited to mammalian vaults. Two independent 2026 preprints showed that prokaryotic MVP homologues can assemble into vault structures. Athukoralage et al. (2026) identified MVP homologues across diverse bacterial and archaeal lineages and demonstrated assembly of several prokaryotic and bacteriophage proteins into vault particles. A bacteriophage-derived MVP formed a 96-subunit vault with a larger internal volume than the human particle, and increasing the number of MVP wall repeats produced still larger particles. Independently, Lachowicz et al. determined bacterial vault structures with barrel and cap symmetries substantially different from those of previously characterized eukaryotic vaults (Lachowicz et al., 2026). Extending loop-based conjugation to such prokaryotic scaffolds would provide alternative particle geometries and larger internal capacities for surface antigen display and cargo loading.

Future work evaluating virus neutralization, secretory IgA in respiratory washes, and B-cell receptor cross-linking across LOW, MEDIUM, and HIGH antigen densities will further define the potential of these engineered particles. Overall, dual-loop DogTag insertion provides a straightforward way to display complex viral antigens at tunable densities on human vault nanoparticles without redesigning the underlying scaffold. For now, our structural and in vivo findings demonstrate the feasibility of the vault platform for high-density covalent surface display of complex viral antigens.

## Materials and methods

### Dual-DogTag MVP construct

The coding sequence of human major vault protein (MVP) was engineered to incorporate two DogTag insertions within selected surface-exposed loops. The first insertion was introduced into repeat domain R3 by replacing the native amino acid residues 132–135 (KDGD) with the DogTag sequence, while preserving the flanking residues D131 and K136. Similarly, the second insertion was introduced into repeat domain R7 by replacing residues 346–347 (EE), while retaining D345 and K348. Consequently, each engineered MVP monomer contained two DogTag sequences, providing a theoretical maximum of 156 DogTag sites per assembled vault particle comprising 78 MVP subunits. The engineered MVP sequence was codon-optimized for expression in human cells, commercially synthesized, and cloned into the BamHI restriction site of the pcDNA3.1(+) expression vector by GenScript (Piscataway, NJ, USA). The resulting plasmid was propagated in NEB Stable Competent *Escherichia coli* cells (New England Biolabs, Ipswich, MA, USA), and plasmid DNA was purified using the NucleoBond Xtra Midi kit (Macherey-Nagel, Düren, Germany) according to the manufacturer’s instructions.

### HA-DogCatcher construct

The H5N1 Clade 2.3.4.4b influenza hemagglutinin (HA)–DogCatcher construct was designed based on the HA sequence corresponding to GenBank accession number YCB31181.1. The soluble HA ectodomain was engineered by replacing the native furin-cleavage site with the GSAS sequence and introducing three stabilizing substitutions (H371W, K396I, and E448I), as previously described (Milder et al., 2022). The HA ectodomain was fused at its C-terminus to a T4 fibritin foldon trimerization domain, followed by a flexible (GGGGS)₃ linker and the DogCatcher sequence. An N-terminal CD33 signal peptide for secretion and a polyhistidine tag for purification were incorporated, as illustrated in Fig. 3A. The human codon-optimized HA–DogCatcher gene was synthesized by GenScript and inserted into pcDNA3.1(+) at the BamHI restriction site. Plasmid amplification in NEB Stable Competent *E. coli* and subsequent purification using the NucleoBond Xtra Midi kit were performed. HA–DogCatcher was expressed independently of the vault scaffold.

### Expression and purification of DogTag-functionalized vault nanoparticles and HA–DogCatcher

Suspension HEK293 cells maintained in Expi293 Expression Medium (Thermo Fisher Scientific, Waltham, MA, USA) were transfected with the dual-DogTag MVP expression plasmid using PEIpro (Polyplus, Illkirch, France) according to the manufacturer’s recommendations. Cells were incubated at 37 °C for four days before harvesting, while mock-transfected cells were processed in parallel as controls. Cell pellets were resuspended in PBS adjusted to a final NaCl concentration of 300 mM and supplemented with SIGMAFAST EDTA-free Protease Inhibitor Cocktail Tablets (Sigma-Aldrich, St. Louis, MO, USA) and 0.05% Triton X-100. Cell lysis was performed through five freeze–thaw cycles, followed by treatment with Denarase (250 U/mL) for 2 h at room temperature. The resulting lysates were clarified by centrifugation at 15,000 × g for 15 min to remove cellular debris, and the collected supernatants were passed through a 0.2-µm membrane filter. A second Denarase treatment was subsequently performed before purification by Capto Core 700 chromatography using an ÄKTA go chromatography system (Cytiva, Uppsala, Sweden). The columns were equilibrated with PBS containing a final NaCl concentration of 300 mM, and the flow-through containing vault particles was collected for further purification. Size-exclusion chromatography was performed using qEV1 35-nm columns (Izon Science, Christchurch, New Zealand), equilibrated with the same buffer. For each run, 1 mL of sample was loaded onto the column, followed by 3 mL of equilibration buffer. Four consecutive particle-containing fractions (F5–F8), each corresponding to 700 µL, were collected, with the highest particle abundance consistently observed in fractions F6 and F7. Mock-transfected samples underwent the same purification procedure.

HA–DogCatcher was expressed separately in suspension HEK293 cells using the same Expi293 medium and PEIpro-mediated transfection protocol described above, followed by incubation at 37 °C for four days. The HA–DogCatcher-containing culture supernatant was harvested and treated with Denarase prior to purification by immobilized metal affinity chromatography using 5 ml HisTrap FF columns connected to an ÄKTA go system (Cytiva). The columns were equilibrated with PBS containing 300 mM NaCl and 25 mM imidazole, and the samples were supplemented with imidazole to a final concentration of 25 mM before loading. Following sample application, the columns were washed with an equilibration buffer to remove unbound proteins. Bound HA–DogCatcher was subsequently eluted using PBS containing 300 mM NaCl and 300 mM imidazole. The collected protein fractions were desalted using HiPrep columns packed with Sephadex G-25 resin (Cytiva), equilibrated with PBS containing a final NaCl concentration of 300 mM. The concentration of purified HA–DogCatcher was determined using the Pierce Micro BCA Protein Assay Kit (Thermo Fisher Scientific) according to the manufacturer’s instructions.

### SDS-PAGE, TRPS and Scanning electron microscopy

Protein profiles of samples collected at different stages of vault production and purification were examined by SDS-PAGE. Samples were mixed with 6× reducing Laemmli sample buffer, heated at 95 °C for 5 min, and separated on 4–20% Mini-PROTEAN TGX Stain-Free precast gels (Bio-Rad, Hercules, CA, USA) using the Mini-PROTEAN Tetra Cell electrophoresis system. Protein molecular weights were estimated using PageRuler Prestained Protein Ladder (Thermo Fisher Scientific, Waltham, MA, USA). The engineered MVP was detected at an apparent molecular weight of approximately 120 kDa.

Particle concentration and size distribution of qEV-purified vault preparations were assessed by tunable resistive pulse sensing (TRPS) using the Exoid system (Izon Science, Christchurch, New Zealand). The instrument was calibrated with standardized polystyrene nanoparticles, and measurements were conducted using the manufacturer’s recommended consumables and procedures. Samples were diluted as required to achieve suitable particle detection conditions, and data were acquired and processed using Izon Control Suite software. Since TRPS detects particles based on changes in electrical resistance during nanopore translocation, the resulting particle concentrations represent total detectable particles within the purified preparations rather than exclusively MVP-containing vaults. For particle-normalized HA-loading calculations, the particle count was restricted to the dominant vault-sized region of the TRPS distribution, comprising the three major adjacent peaks corresponding to the expected vault-sized population; larger-particle populations were excluded

For SEM, a 10 µL aliquot of the vault nanoparticle suspension was deposited onto glass coverslips and allowed to air-dry at room temperature for 30 min. Following drying, the samples were sputter-coated with a 10 nm-thick gold layer using a Quorum Q150RS sputter coater to provide electrical conductivity. The samples were subsequently examined at an accelerating voltage of 2 kV using a ZEISS Sigma 500 field-emission scanning electron microscope (FE-SEM) equipped with an SE2 (secondary electron) detector at the Electron Microscopy Laboratory of the Izmir Biomedicine and Genome Center (IBG), Türkiye.

### HA–DogCatcher conjugation and anti-MVP immunoblotting

Purified HA–DogCatcher protein was mixed with dual-DogTag vault nanoparticles and incubated at room temperature for 15 min to promote spontaneous covalent conjugation between DogTag and DogCatcher. HA–SpyTag protein was included as a non-cognate control to assess conjugation specificity. Covalent attachment of HA–DogCatcher to MVP was evaluated by Western blotting under reducing conditions. Following SDS-PAGE, proteins were transferred onto an Amersham nitrocellulose membrane (Cytiva, Uppsala, Sweden) at 250 mA for 45 min. Membranes were blocked with 5% bovine serum albumin (BSA) and subsequently incubated with a rabbit polyclonal anti-MVP antibody (A305-172A; Thermo Fisher Scientific, Waltham, MA, USA), followed by an HRP-conjugated anti-rabbit secondary antibody. Immunoreactive proteins were detected using Amersham enhanced chemiluminescence (ECL) reagents (Cytiva). Under reducing conditions, higher-molecular-weight MVP-reactive bands were interpreted as consistent with individual MVP polypeptides covalently conjugated to one or two HA–DogCatcher chains, rather than intact HA trimers.

### Controlled HA-loading series

To examine the effect of antigen input on vault loading, a fixed amount of vault material was incubated with increasing amounts of HA-DogCatcher. The loading series contained 0, 25, 50, 100 and 200 µg HA-DogCatcher. After conjugation, samples were subjected to qEV size-exclusion chromatography to recover particle-associated material and reduce unbound HA.

A separate maximal-loading preparation was produced by adding 800 µg HA-DogCatcher to lyophilized vault material. Because this preparation differed in volume and handling from the 0–200 µg loading series, it was analysed as a separate high-loading preparation rather than as an additional point in the linear loading series.

Reaction extent was assessed from the progressive loss of the unmodified MVP band on stain-free SDS–PAGE together with the appearance of higher-molecular-weight conjugated species. Under reducing conditions, these shifted species correspond to individual MVP chains carrying one or two covalently attached HA–DogCatcher polypeptide chains and were therefore used as evidence of DogTag reaction rather than as a direct measure of intact HA trimers per vault. Purified HA–DogCatcher was additionally loaded at 0.5, 1.0 and 2.0 µg per lane as a same-gel reference series for the migration and stain-free response of the unconjugated antigen.

### Estimation of reacted DogTag sites by gel densitometry

The intensities of the unmodified MVP band and the two bands corresponding to MVP carrying one or two HA–DogCatcher chains were measured from stain-free gel images after background subtraction.

The percentage decrease in the intensity of the unmodified MVP band compared with the vault-only control was used to estimate the proportion of MVP subunits that had reacted with HA–DogCatcher. This percentage was used to calculate the number of modified MVP subunits per vault, assuming 78 MVP subunits per particle and comparable amounts of MVP in each sample.

Since stain-free fluorescence depends on the tryptophan content of proteins, the intensities of the bands corresponding to MVP carrying one or two HA–DogCatcher chains were divided by their respective numbers of tryptophan residues (18 and 29). The relative intensities of these two bands after correction were used to estimate how many of the modified MVP subunits carried one or two HA–DogCatcher chains. The total number of reacted DogTag sites per vault was then calculated by counting one site for each singly modified MVP and two sites for each doubly modified MVP.

Purified HA–DogCatcher (0.5, 1 and 2 µg) was run on the same gel to examine the relationship between protein amount and band intensity. The bands in the 25 µg HA preparation were too faint for reliable quantification. For the 800 µg preparation, which was prepared under different conditions, only the decrease in the unmodified MVP band was evaluated.

The calculations are provided in Supplementary Table 1A and estimate the number of reacted DogTag sites rather than the number of attached HA trimers.

### Estimation of HA trimers per particle

HA loading was estimated from the amount of HA recovered in the qEV particle fraction together with the corresponding TRPS-derived vault-sized particle count. The TRPS denominator was calculated from the dominant vault-sized region of the particle-size distribution, comprising the three major adjacent peaks, while larger-particle populations were excluded. For the 25–200 µg loading series, pooled qEV volumes were approximately 0.700–0.750 mL and the corrected vault-sized particle recovery was approximately 3.5 × 10¹¹ particles per pooled preparation.

For the separate maximal-loading preparation, HA was quantified by ELISA using the identical HA–DogCatcher protein as the calibration standard. The preparation contained approximately 44–67 µg HA associated with approximately 9 × 10¹¹ TRPS-defined vault-sized particles.

A working molecular mass of approximately 100 kDa per HA–DogCatcher protomer and 300 kDa per trimer was used to convert HA mass into trimer equivalents. The resulting values are preparation-level particle-associated HA trimer-equivalent estimates. They do not by themselves constitute a direct measurement of the number of covalently attached intact HA trimers on each individual vault.

The concentration of HA in qEV-purified vault preparations was determined by enzyme-linked immunosorbent assay (ELISA) using an HRP-conjugated anti-His-tag antibody. Briefly, purified HA–DogCatcher protein was used to generate a standard curve through twofold serial dilutions ranging from 8 to 0.0078 µg/mL, with 11 standard concentrations analysed in duplicate. Vault–HA samples were diluted 1:50, 1:100, and 1:200, and 50 µL of each dilution was added to 96-well microplates alongside the HA standards. Undecorated vault preparations served as negative controls, while purified HA–DogCatcher was included as a system suitability control. Blank wells were included to determine background absorbance. Plates were incubated for 2 h at room temperature to allow protein adsorption, followed by blocking with 5% bovine serum albumin (BSA) for 1 h at room temperature. After washing, an HRP-conjugated 6×His-tag monoclonal antibody (Proteintech, Rosemont, IL, USA) was added and incubated for 1 h at room temperature. Following additional washing, 3,3′,5,5′-tetramethylbenzidine (TMB) substrate was added for colorimetric detection. The reaction was terminated by the addition of sulfuric acid (H₂SO₄), and absorbance was measured at 450 nm. A four-parameter logistic (4PL) regression model was fitted to the HA standard curve, and HA concentrations in the purified vault preparations were calculated by interpolation from the fitted curve, accounting for the corresponding sample dilution factors.

For the separate high-loading preparation, total HA was measured using the same HA-DogCatcher protein as the ELISA standard. The high-loading preparation contained approximately 44–67 µg HA associated with a vault-sized TRPS population of approximately 9 × 10¹¹ particles. A working molecular mass of 100 kDa per HA-DogCatcher protomer and 300 kDa per HA trimer was used for conversion of HA mass to trimer number. The average number of HA trimers per particle was calculated as:

HA trimers per particle = [(HA concentration × pooled volume × 10^-6) / particle number] × [6.02214076 × 10^23 / 3.0 × 10^5]

where HA concentration is in µg/mL, pooled volume is in mL and 3.0 × 10^5 g/mol is the working molecular mass of one HA-DogCatcher trimer.

For the high-loading preparation, measured total HA mass was substituted directly for HA concentration × pooled volume. Using the corrected ∼3.5 × 10¹¹-particle denominator for the controlled series, the +50, +100 and +200 µg preparations corresponded to 30.9–33.1, 62.2–66.7 and 73.1–78.3 HA trimer equivalents per particle, respectively. These calculations defined the approximate LOW (∼32), MEDIUM (∼64) and HIGH (∼124) preparation-level loading states used for structural modelling; HIGH was based on the independent maximal-load preparation (98.1–149.4; midpoint, 123.8). The 149-trimer value obtained from the upper end of the high-loading mass estimate was treated only as an upper-bound stress-test value and was not used as a main structural state.

These values represent average estimates for each preparation and do not indicate the exact number of HA molecules on an individual particle.

### Negative-stain STEM

Empty and highly HA-loaded vault nanoparticles were examined by negative-stain scanning transmission electron microscopy (STEM). Briefly, a 10 µL aliquot of the vault nanoparticle suspension was deposited onto the carbon-coated face of Formvar/carbon-coated copper grids and allowed to adsorb for 1 min. The grids were then briefly rinsed three times with ultrapure water to remove unbound material, followed by negative staining with 2% (w/v) uranyl acetate (prepared in distilled water) for 1 min. Excess staining solution was blotted with filter paper, and the grids were air-dried at room temperature for 30 min. Imaging was performed in STEM mode at an accelerating voltage of 8 kV using a ZEISS Sigma 500 field-emission scanning electron microscope (FE-SEM) equipped with a STEM detector at the Electron Microscopy Laboratory of the Izmir Biomedicine and Genome Center (IBG), Türkiye.

### Particle-level analysis of near-end-on STEM images

Four isolated near-end-on particles from the highly loaded preparation were selected as the high-confidence primary set (1916_P1, 1929_P1, 1933_P1 and 1935_P1). Two additional particles were retained as lower-confidence sensitivity examples but were not included in the primary comparison.

For each particle, the particle centre was defined manually and analysis was restricted to a radial region 8–45 nm from the centre. Image intensity was normalized using a robust local-contrast procedure within each particle region. Candidate peripheral densities were detected with an apparent diameter constraint of 7–10 nm. The primary detection threshold was 0.24. Detector sensitivity was assessed across thresholds from 0.20 to 0.28.

Nearest-neighbour distances were calculated separately within each particle. Distances between detections belonging to different particles were never included. For each particle, the number of detected peripheral densities and the median nearest-neighbour distance were recorded. The four particles were treated as individual image-level observations rather than independent biological replicates.

### Structural modelling of HA-loaded vaults

The structure of the human vault (PDB 9QW9), containing 78 MVP subunits, was used to determine the positions of the DogTag attachment sites. These sites were defined as the midpoints between the central carbon (Cα) atoms of D131 and K136 in R3 and D345 and K348 in R7. Since each MVP subunit carries two DogTags, there were 156 potential attachment sites on the vault surface.

The structure of the HA–DogCatcher trimer was obtained using AlphaFold. In this structure, residues 1–534 correspond to HA, residues 535–561 to the foldon domain, residues 562–576 to the linker and residues 577–680 to DogCatcher. During modelling, the shape of the HA–foldon region (residues 14–561) was kept unchanged. Since the DogTag, DogCatcher and linker regions allow some flexibility in the attachment of HA to the vault, their exact conformations were not modelled. Instead, we assumed a distance of 4.5 nm between the attachment site on MVP and the foldon domain and allowed HA trimers to tilt by up to 45° from the outward-facing direction.

### Arrangement of HA trimers on the vault surface

Based on the experimentally estimated HA loading levels, we generated three models containing 32, 64 and 124 HA trimers per vault. To reduce the number of calculations required when arranging HA trimers on the vault surface, we approximated the shape and dimensions of each HA trimer using 27 overlapping spheres derived from its atomic structure. Once suitable arrangements were obtained, the original atomic coordinates were used to check for steric clashes.

Different combinations of attachment sites and HA orientations were tested to find arrangements in which HA trimers did not overlap with one another or with the vault surface. A minimum separation of 0.15 nm from the vault surface was maintained. To obtain the model containing 124 HA trimers, we started with an arrangement containing 107 trimers and adjusted their positions and orientations to make room for additional trimers. The same criteria were used for all three models. The three structural models were generated using Python scripts and visualized using UCSF ChimeraX (version 1.12).

### Measurement of distances between HA trimers

To confirm that the HA trimers could be accommodated in the proposed arrangements, we examined their atomic coordinates for steric clashes. Distances of less than 2 Å between atoms or overlaps of more than 0.6 Å between their van der Waals radii were considered steric clashes. No such clashes were found between the HA–foldon structures or between these structures and the vault surface in the final models.

The centre of each HA trimer was determined by averaging the coordinates of its alpha-carbon (Cα) atoms. The three-dimensional distance between the centre of each HA trimer and that of its closest neighbouring trimer was calculated using Python scripts. The minimum, maximum, mean and median distances were determined for each of the three loading levels (LOW, MEDIUM and HIGH).

To determine whether these distances depended on the assumed tilt angle of HA, the models were also generated using maximum tilt angles of 30° and 60° instead of 45°, without changing the other parameters. Structural models were visualized using UCSF ChimeraX (version 1.12).

### Comparison of the HIGH model with STEM images

Since most of the highly loaded vaults appeared end-on in the STEM images, we examined how the model containing 124 HA trimers would appear when viewed end-on. Using Python scripts, the centres of all 124 HA trimers were projected onto a two-dimensional plane perpendicular to the long axis of the vault. Additional projections were generated at tilt angles of 5°, 10° and 15°. For each tilt angle, 12 views were generated at 30° intervals around the long axis of the vault. Together with the direct end-on projection, this provided 37 simulated views. To approximate the appearance of HA in the STEM images, each projected HA centre was represented by a blurred spot using a Gaussian function with a full width at half maximum of 10 nm. This allowed closely positioned HA trimers to appear as a single spot. The number of distinct spots and the median nearest-neighbour distance between them were calculated for each simulated image. These values were compared with the corresponding measurements obtained from four isolated, end-on vault particles in the STEM images.

### Mouse immunization

Female BALB/c mice aged 6–8 weeks were used under institutional animal ethics approval. Mice received a subcutaneous prime on day 0 followed by an intranasal boost on day 28, and serum was collected on day 38. Antigen-containing groups received 2 µg total HA at prime and 1 µg total HA at boost. Vault-containing groups received approximately 3.3 × 10^10^ particles at prime and 1.65 × 10^10^ particles at boost.

Experimental groups were vault only, free HA, vault mixed with non-conjugated HA, HA-decorated vault and HA plus 50 µg CpG ODN as an adjuvanted positive-control.

All animal procedures were conducted in accordance with protocols approved by the Izmir Biomedicine and Genome Center Animal Experiments Local Ethics Committee (IBG-HADYEK; Protocol No. 2024-016).

### Assessment of HA and Vault-specific serum antibody responses and statistical analysis

HA-specific serum IgA, IgG, IgG1, and IgG2a antibody responses and vault specific IgG responses were determined by enzyme-linked immunosorbent assay (ELISA). Briefly, 96-well microplates were coated with purified recombinant HA-SpyTag protein (1 µg/mL, 50 µL/well) or 2.5 x 10^9^ Vault particles/well and incubated overnight at 4 °C. The following day, plates were blocked with 5% bovine serum albumin (BSA) in PBS containing Tween-20 (PBST) for 2 h at room temperature. Following washing, serum samples were initially diluted 1:50 and subsequently subjected to fourfold serial dilutions to generate eight dilutions ranging from 1:50 to 1:819,200. Diluted sera were transferred to HA or vault-coated plates and incubated overnight at 4 °C. After washing, alkaline phosphatase (AP)-conjugated anti-mouse IgA, IgG, IgG1, or IgG2a secondary antibodies (SouthernBiotech, Birmingham, AL, USA) were added at a 1:1,000 dilution and incubated for 2 h at room temperature. Plates were subsequently washed and developed using p-nitrophenyl phosphate (pNPP) substrate (Thermo Fisher Scientific, Waltham, MA, USA). Absorbance was measured at 405 nm using a Varioskan LUX multimode microplate reader (Thermo Fisher Scientific). The absorbance cutoff for each assay was calculated as the mean optical density of placebo-control samples plus four standard deviations (mean + 4 SD). Antibody endpoint titres were defined as the reciprocal of the highest serum dilution yielding an absorbance value above the corresponding cutoff.

Statistical analyses were performed using GraphPad Prism software (GraphPad Software, Boston, MA, USA). Individual mice were considered independent biological replicates, and antibody endpoint titres were log10-transformed prior to analysis. Comparisons among experimental groups were performed using Šídák’s multiple comparisons test. Differences were considered statistically significant at p < 0.05.

