## Supplementary material for "Dual-loop DogTag engineering enables modular antigen display on human vault nanoparticles": FigureS1

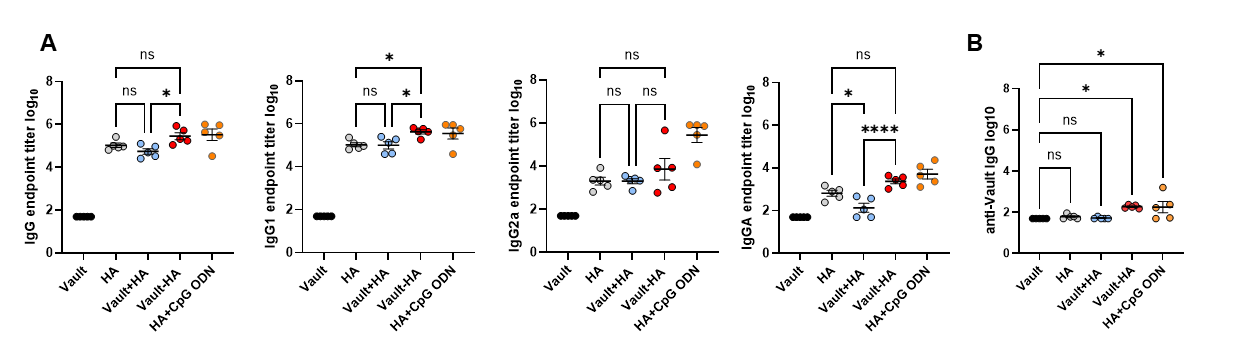

**Figure S1. Humoral immune responses induced by HA-decorated vaults in mice.** Mice (n= 5 per group), were primed subcutaneously (2 µg HA) and boosted intranasally (1 µg HA) with empty vaults (Vault), soluble HA (HA), a physical mixture of empty vaults and HA (Vault+HA), HA-decorated vaults (Vault-HA), or HA adjuvanted with CpG ODN (HA+CpG ODN). Serum samples collected 10 days after boosting were analyzed by ELISA. (A) HA-specific serum total IgG, IgG1, IgG2a, and IgA log_10_ endpoint titers are shown. (B) Anti-vault serum IgG log_10_ endpoint titers measured on vault-coated plates. Statistical significance was determined by one-way ANOVA followed by Šídák's multiple-comparisons test (*p < 0.05,**p < 0.01, ***p < 0.001, ****p < 0.0001; ns, not significant)
