## Supplementary Information for "Dual-loop DogTag engineering enables modular antigen display on human vault nanoparticles"

**Supplementary information: Structural modelling of HA-loaded vaults**

**Supplementary methods**

**Structure of the vault and HA trimer** The human vault structure (PDB 9QW9) contains 78 MVP subunits. On each subunit, the two potential HA attachment points were defined as the midpoints between the backbone carbon (Cα) atoms of D131 and K136 in repeat R3, and of D345 and K348 in repeat R7. This gave 156 potential attachment sites arranged in four rings around the vault body.

The top-ranked AlphaFold model of the HA–DogCatcher trimer was used (residue ranges are listed in Table S2). The main HA–foldon body (residues 14–561) was kept fixed as a rigid shape, while the flexible N-terminal tail (residues 1–13) was left out. Because DogTag, DogCatcher and the linker can bend and move, their exact atomic positions were not calculated. Instead, we assumed a 4.5 nm spacing from the attachment site on the vault to the foldon domain, and allowed each HA trimer to tilt by up to 45° away from straight out. These values were modelling assumptions rather than direct measurements.

**Arrangement of HA trimers on the vault surface** Using Python scripts, we built models with 32, 64 and 124 HA trimers per vault to match the experimentally estimated HA loading levels. To speed up the calculations, the shape of each HA–foldon trimer was represented by 27 overlapping spheres based on its atomic positions. The script tested different attachment sites and tilt angles while keeping at least 0.15 nm of empty space between each HA trimer and the vault surface. Trimers were placed automatically rather than by hand, and they were not forced into equal numbers on each ring. To reach 124 trimers for the HIGH model, we started from a 107-trimer arrangement and adjusted existing trimers to make room for additional ones. All three loading levels used the same geometric rules. Molecular images were made in UCSF ChimeraX (version 1.12).

**Checking the models and calculating HA spacing** Once the trimers were placed, we checked the full atomic coordinates of the HA–foldon structures to make sure nothing overlapped physically. Any pair of atoms closer than 2 Å, or with overlapping atomic radii (van der Waals overlap) greater than 0.6 Å, was counted as a clash.

We found the centre of each HA trimer by averaging the positions of its Cα atoms, then used Python to measure the straight-line 3D distance from each trimer centre to its closest neighbour. From these values, we recorded the minimum, maximum, mean and median distances.

To check whether the assumed 45° tilt limit affected spacing, we repeated the modelling with maximum tilt angles of 30° and 60° while keeping all other settings unchanged. All nine models reached their target trimer counts with zero physical clashes (Fig. S3 and Table S3).

**Comparison with STEM images** We defined the long axis of the vault from the average positions of the top and bottom rings of attachment sites, then used Python to project the centres of all 124 HA trimers onto a flat plane perpendicular to that axis (an end-on view). We also generated tilted views at 5°, 10° and 15°, rotating the vault in 30° steps (12 views per tilt angle), giving 37 simulated views in total.

To mimic how the particles look at the resolution of negative-stain STEM, each projected HA centre was blurred into a 10 nm wide circular spot (Gaussian width at half-maximum). This 10 nm blur was chosen for the simulation rather than measured from the microscope. Overlapping spots merged together, and each bright peak in the resulting image was counted as one visible spot. We compared the number of visible spots and the median distance between neighbouring spots against measurements from four isolated, nearly end-on vault particles in the STEM images. This comparison was descriptive only and did not establish the actual number or layout of HA trimers on individual particles.

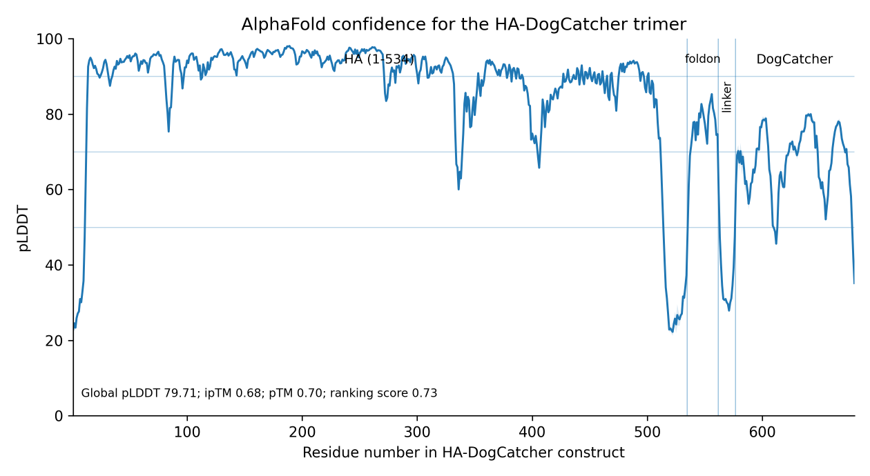

**Supplementary Figure S2. Confidence of the AlphaFold HA–DogCatcher structure** AlphaFold per-residue confidence scores (pLDDT) across the HA–DogCatcher sequence, averaged over the three protein chains (shading shows the range across chains). Confidence is high across most of the HA region and lower in the flexible linker. Overall model scores were pLDDT = 79.71, overall fold confidence (pTM) = 0.70, subunit-interface confidence (ipTM) = 0.68, and ranking score = 0.73. Higher pLDDT values indicate higher confidence in the predicted local structure.

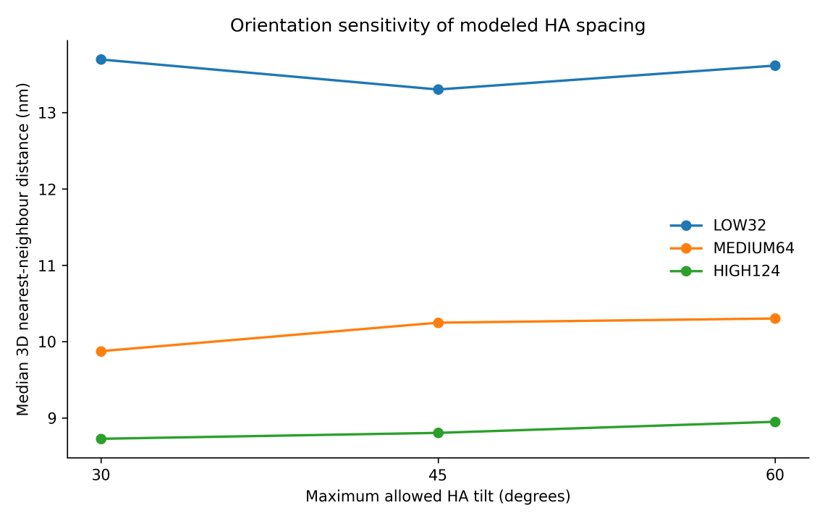

**Supplementary Figure S3. Effect of HA tilt angle on the distance between trimers** Median 3D nearest-neighbour distances between HA trimers in the LOW (32), MEDIUM (64) and HIGH (124) models at maximum tilt angles of 30°, 45° and 60°. At every tilt angle, trimers were spaced closer together as HA loading increased.

**Supplementary Table S2. Parameters used for structural modelling**

| **Parameter** | **Value or definition** |
| --- | --- |
| Vault structure | PDB 9QW9 (78 MVP subunits) |
| R3 attachment site | Midpoint between Cα atoms of D131 and K136 |
| R7 attachment site | Midpoint between Cα atoms of D345 and K348 |
| Potential attachment sites | 156 (two per MVP subunit, arranged in four rings) |
| HA–DogCatcher structure | Highest-ranked AlphaFold trimer model |
| Residues in the HA–DogCatcher model | HA: 1–534; foldon: 535–561; linker: 562–576; DogCatcher: 577–680 |
| Region kept rigid | Residues 14–561 (HA–foldon) |
| Attachment site to foldon distance | 4.5 nm (assumed) |
| Maximum HA tilt | 45° in the main models; also tested at 30° and 60° |
| Clearance at the vault surface | 0.15 nm |
| Approximation of the HA shape | 27 overlapping spheres |
| HA trimers per model | LOW: 32; MEDIUM: 64; HIGH: 124 |
| Criteria for steric clashes | Interatomic distance <2 Å or van der Waals overlap >0.6 Å |
| Simulated views of the HIGH model | 37 (0°, 5°, 10°, 15° tilts; 30° intervals at each non-zero tilt) |
| Width of simulated spots | 10 nm (full width at half maximum; assumed) |

The AlphaFold model used was fold_ha_with_dogcatcher_model_0.cif.

**Supplementary Table S3. Effect of HA tilt angle on the distance between trimers**

| **Maximum HA tilt** | **LOW (32)** | **MEDIUM (64)** | **HIGH (124)** | **Sites suitable for placement** |
| --- | --- | --- | --- | --- |
| 30° | 13.70 (9.50–22.84) | 9.87 (8.35–16.68) | 8.73 (7.65–11.43) | 152/156 |
| 45° | 13.30 (8.74–23.53) | 10.25 (8.35–15.96) | 8.80 (7.86–10.65) | 156/156 |
| 60° | 13.62 (8.84–21.89) | 10.30 (8.26–16.64) | 8.95 (7.97–11.82) | 156/156 |

Values are median nearest-neighbour distances (nm), with minimum and maximum distances in parentheses (these show the range within each model, not confidence intervals). The last column shows how many of the 156 attachment sites could physically fit an HA trimer under each tilt limit, not how many were occupied. No physical clashes were detected in any of the nine final models.
