## Supplementary Table 1 for "Dual-loop DogTag engineering enables modular antigen display on human vault nanoparticles"

**Supplementary Table 1. Quantification of HA loading in the Figure 3C–D vault preparations**

**A. Stain-Free gel analysis of HA–DogCatcher conjugation**

**A1. Loss of unmodified MVP across the loading series**

| **Lane** | **Preparation** | **Unmodified MVP signal, S0 (AU)** | **MVP remaining (%)** | **Loss of unmodified MVP (%)** | **Modified MVPs/vault, Nmod** | **Control** |
| --- | --- | --- | --- | --- | --- | --- |
| 1 | Vault only | 2,913,824 | 100.0 | 0.0 | 0.00 | reference |
| 2 | +25 µg | 2,817,040 | 96.7 | 3.3 | 2.59 | low-signal |
| 3 | +50 µg | 2,120,508 | 72.8 | 27.2 | 21.24 | LOW |
| 4 | +100 µg | 1,618,542 | 55.5 | 44.5 | 34.67 | MEDIUM |
| 5 | +200 µg | 956,124 | 32.8 | 67.2 | 52.41 | HIGH |
| 6 | +800 µg | 497,444 | 17.1 | 82.9 | 64.68* | separate maximal-load preparation |

For each lane, the densitometric intensity of the unmodified MVP band was normalized to that of the vault-only control (2,913,824 AU) to obtain the percentage of residual unmodified MVP, with the complementary depleted percentage representing MVP shifted into higher-molecular-weight conjugates. Multiplying this shifted fraction by the canonical stoichiometry of 78 MVP subunits per vault yielded the mean number of modified MVP subunits per vault. *The +800 µg preparation was generated under distinct reaction conditions; its apparent value is shown for descriptive comparison only and was excluded from quantitative site-occupancy analysis.

**A2. Shifted-band measurements, tryptophan correction and reacted-site calculation**

| **Preparation** | **MVP+1HA, S1 (AU)** | **MVP+2HA, S2 (AU)** | **Raw S2/S1** | **q1 = S1/18** | **q2 = S2/29** | **Trp-corrected q2/q1** |
| --- | --- | --- | --- | --- | --- | --- |
| **+25 µg** | ~271 (trace) | ND | — | ~15.06 | — | — |
| **+50 µg** | ~1,659 | ~2,990 | 1.80 | 92.17 | 103.10 | 1.119 |
| **+100 µg** | ~2,485 | ~4,789 | 1.93 | 138.06 | 165.14 | 1.196 |
| **+200 µg** | 2,963 | 5,811 | 1.96 | 164.61 | 200.38 | 1.217 |
| **+800 µg** | present; not quantified | present; not quantified | — | — | — | — |

Densitometric intensities of the shifted bands corresponding to MVP+1HA and MVP+2HA were quantified from the Stain-Free gel TIFF image using narrow, local-background-corrected regions of interest. Because Stain-Free fluorescence scales proportionally with tryptophan (Trp) content-and engineered MVP and HA–DogCatcher contain 7 and 11 Trp residues, respectively-the MVP+1HA and MVP+2HA band intensities were normalized to their respective total Trp counts (18 and 29 residues) to obtain relative molar abundances and the Trp-corrected molar ratio of MVP+2HA to MVP+1HA. The shifted-band signal in the +25 µg lane was too weak for reliable quantification, and the +800 µg lane was excluded as a separately prepared maximal-load sample.

| **Preparation** | **Nmod = N1+N2** | **N1 (MVP+1)** | **N2 (MVP+2)** | **Reacted sites, C=N1+2N2** | **Rounded estimate** | **Strict bound from Nmod** |
| --- | --- | --- | --- | --- | --- | --- |
| **+50 µg** | 21.24 | 10.02 | 11.21 | 32.45 | ~32 | 21.24–42.47 |
| **+100 µg** | 34.67 | 15.79 | 18.89 | 53.56 | ~54 | 34.67–69.35 |
| **+200 µg** | 52.41 | 23.63 | 28.77 | 81.18 | ~81 | 52.41–104.81 |

The modified MVP population (from A1) was apportioned into MVP+1HA and MVP+2HA subunits according to their respective Trp-corrected molar fractions. The total number of reacted DogTag sites per vault-equivalent was then calculated by counting one reacted site per MVP+1HA subunit and two per MVP+2HA subunit. Strict theoretical bounds span the limiting cases in which all modified MVP subunits carry either one or two attachments. These values estimate polypeptide-level DogTag engagement rather than the number of intact HA trimers per individual vault.**A3. Same-gel HA–DogCatcher standards**

| **Purified HA–DogCatcher loaded** | **Integrated Stain-Free signal (AU)** | **Median-local-background sensitivity value (AU)** |
| --- | --- | --- |
| **0.5 µg** | 23,493 | 25,009 |
| **1.0 µg** | 44,970 | 49,695 |
| **2.0 µg** | 80,158 | 90,072 |

Purified HA–DogCatcher standards (0.5, 1.0, and 2.0 µg) were run on the same gel and integrated using both primary and median-local-background corrections, showing linear Stain-Free response across this range (R² = 0.997 for both analyses). These standards served as a same-gel linearity and non-saturation controls; the reacted-site calculation in A2 is based solely on internal MVP band ratios.

**B. Figure 3D qEV ELISA/TRPS analysis**

| **Preparation** | **Post-qEV HA** | **HA recovered** | **Vault-sized particles** | **Estimated HA trimers/particle** | **Role** |
| --- | --- | --- | --- | --- | --- |
| **+25 µg** | 9.5 µg/mL | 6.65–7.13 µg | ~3.5 × 10¹¹ | 38.1–40.9 | context |
| **+50 µg** | 7.7 µg/mL | 5.39–5.78 µg | ~3.5 × 10¹¹ | 30.9–33.1 | LOW ~32 |
| **+100 µg** | 15.5 µg/mL | 10.85–11.63 µg | ~3.5 × 10¹¹ | 62.2–66.7 | MEDIUM ~64 |
| **+200 µg** | 18.2 µg/mL | 12.74–13.65 µg | ~3.5 × 10¹¹ | 73.1–78.3 | high-loading context |
| **+800 µg** | — | 44–67 µg | ~9 × 10¹¹ | 98.1–149.4 | HIGH ~124 |

For the +25 to +200 µg loading series, post-qEV ELISA concentrations were multiplied by the pooled qEV fraction volume (0.70–0.75 mL) to determine total recovered HA mass, whereas total recovered HA (44–67 µg) was measured directly for the separate +800 µg preparation. Recovered HA mass was converted into trimer molecule counts using a working trimer mass of 300 kDa per HA–DogCatcher trimer, and then divided by the TRPS-derived vault-sized particle count (~3.5 × 10¹¹ for the titration series; ~9 × 10¹¹ for the +800 µg preparation) to estimate HA trimer equivalents per particle.
